# From video-derived feeding behaviour to cow-level nutritional deviation signals: A dairy digital-twin decision-support framework

**DOI:** 10.64898/2026.08.14.744981

**Authors:** Shreya Rao, Suresh Raja Neethirajan

## Abstract

Continuous video offers a dynamic view of dairy-cow behaviour, but its value for precision nutrition depends on alignment with physiological context. We developed a dairy digital-twin framework that fuses identity-associated behavioural records from an established video-analytics layer with body weight, milk production, milk fat, parity and days in milk. The 16-cow analytical cohort was monitored for up to 11 days in one tie-stall barn, yielding 153 cow-days after quality exclusions applied before model fitting. NRC (2001) expected dry matter intake provided a transparent physiological reference compatible with the daily records. In matched leave-one-cow-out analysis, adding video-derived feeding duration to body weight, fat-corrected milk and days in milk reduced RMSE from 2.015 to 1.763 kg DM/day, MAE from 1.267 to 1.159 kg DM/day and MAPE from 4.5% to 4.2%, while R² increased from 0.500 to 0.617. A secondary reduced index achieved RMSE 2.362 kg DM/day and MAPE 5.8% across held-out cows. Cow-level analysis delineated the operating domain: median per-cow MAPE was 4.15%, whereas the sole cow at 15 days in milk had MAPE 28.8%. Two independently recorded veterinary events were temporally concordant with unusual feeding trajectories, providing descriptive biological context. Because the endpoint was NRC-derived, these metrics quantify reference reconstruction rather than accuracy against observed intake. By converting continuous behavioural events into auditable cow-day states, the framework links physical animals to physiologically contextualized digital counterparts and establishes a scalable foundation for operator-focused dairy digital-twin decision support.

## 1. Introduction

Precision livestock farming now delivers continuous, complementary descriptions of animal behaviour and physiology from cameras, wearable sensors, milking systems and farm databases [1,2]. The next opportunity is to unite these streams at the individual-cow level. A behaviour classifier identifies feeding, lying or rumination, while a nutritional equation describes expected state from lactation stage, body size and milk output. Linking the two reveals whether an observed cow-day is concordant with physiological context while preserving the source, timing and operating domain of the resulting signal. This study develops that integration as an auditable cow-day representation and evaluates it under animal-held-out conditions.

Dry matter intake (DMI) provides a demanding and agriculturally important test of this integration. Intake underpins milk production, metabolic adaptation and feed efficiency, while reflecting the combined influence of body size, lactation stage, ration characteristics and physiological state [3–6]. The transition period is especially consequential because nutrient demand rises more rapidly than voluntary intake, increasing susceptibility to negative energy balance and metabolic disruption [7]. Across lactation, changes in feeding, rumination and lying behaviour can provide complementary information about digestion, comfort, competition, disease and environmental stress [4,8–11]. Behaviour is therefore biologically informative, but it must be interpreted in context. Time at the manger is associated with intake, yet eating rate varies among cows, diets and management conditions; feeding duration cannot be equated directly with kilograms consumed[5,6,9,10].

Individual DMI is most convincingly quantified with instrumented feed bins, load cells or comparable feed-weighing systems [12]. Wearable accelerometers and rumination collars provide scalable behavioural measurements and have been evaluated against direct observation, although their performance depends on device placement, behavioural definitions and farm conditions[13]. Video offers a complementary route. It is non-contact, can monitor several animals and behaviours through shared infrastructure, and can build on fixed cameras already used for barn monitoring [2,14]. Feed weighing and video thus answer different but connected questions: one quantifies consumption, whereas the other describes the behavioural process surrounding it. Integrating behavioural continuity with physiological reference modelling offers a practical path toward decision support in facilities where individual feed-intake instrumentation is not routinely available.

Computer-vision methods for cattle have advanced rapidly. Object detectors localise animals, multi-object trackers associate observations through time, and video models distinguish sustained activities across changes in posture and illumination [14–17]. YOLO-family detectors have been applied successfully in complex barn scenes [15]; ByteTrack retains lower-confidence detections that can reduce fragmented trajectories [16]; and temporal architectures such as SlowFast and TimeSformer represent movement patterns that isolated frames cannot capture [17,18]. Our preceding study reported and evaluated a YOLOv11, ByteTrack and TimeSformer pipeline for seven behaviours in a tie-stall dairy barn [19]. That pipeline produces structured records containing cow identity, behaviour, temporal boundaries, duration and confidence. The present study treats those identity-associated records as its established upstream input. Its distinct analytical starting point is the cow-day: aggregating behaviour with body weight, milk production, milk composition, parity and days in milk; relating that representation to a nutritional reference; and evaluating the resulting signal across entirely held-out animals.

This shift in the unit and purpose of inference defines the central novelty of the work. Much of the computer-vision literature treats detection, tracking or classification performance as the endpoint, whereas nutritional models generally operate on periodic physiological and production measurements. The present framework joins these domains through an explicit and auditable cow-day state. It asks whether a separately sensed behavioural variable improves reconstruction of expected nutritional state beyond the physiological variables already represented. Identity association is essential to that question. Herd-level activity summaries can obscure a substantial departure in one animal, whereas a cow-day record preserves the relationship among feeding duration, lactation stage, body weight and milk production. The resulting output is therefore more than an activity label. It is a provenance-aware, cow-level deviation signal that can direct human attention while retaining the source, timing and interpretation of each input.

Nutritional models provide the physiological counterpart to the behavioural stream. NRC (2001) predicts expected DMI from fat-corrected milk, metabolic body weight and an early-lactation adjustment [3], while mechanistic systems such as the Cornell Net Carbohydrate and Protein System organise animal, ration and production variables for precision feeding [20]. NASEM (2021) extends this framework with additional biological information, including parity-related and body-condition effects [21]. In the present records, body-condition score was available at only two time points. NRC (2001) therefore supplied the temporally complete reference compatible with the daily observations. Because body weight, fat-corrected milk and days in milk construct that reference and can also enter the comparison models, much of the agreement is necessarily structural. The informative test is the matched, animal-held-out change obtained when independently sensed feeding duration is added after those physiological inputs are represented. Accordingly, model errors are interpreted as NRC-reference reconstruction, with biological intake accuracy reserved for subsequent feed-weighing validation. Positive and negative deviations denote the direction of model-to-reference disagreement and are intended to support contextual review.

This post-perception integration is naturally situated within a dairy digital twin. Digital twins maintain correspondence between physical entities and computational representations, combine observations with models and create a governed pathway from interpretation to action [22–25]. In livestock systems, staged development is especially valuable because animal responses depend on health, diet, social context and physiological history. The framework presented here implements physical-to-digital updating from video and farm records, constructs a daily per-cow representation, compares it with a nutritional reference and specifies an operator-review gate. We position this contribution at the state-and-interpretation stage of the dairy digital twin: an auditable foundation that creates a direct, testable route to feed-intake calibration, real-time integration and prospective response evaluation.

The study extends a coherent sequence of dairy digital-twin research. Earlier contributions articulated a livestock digital-twin concept [26], synthesised computational architectures for precision dairy nutrition [27] and established the upstream video-perception layer [19]. A preliminary conference communication briefly illustrated the wider architecture over a 24-hour period [28]. The present article advances substantially beyond those foundations through 153 cow-days, explicit cow-day state construction, matched reference-model ablation, leave-one-cow-out evaluation, animal-level operating-domain analysis and a maturity-accounted decision workflow. The novelty resides in this traceable analytical chain from identity-associated behaviour to individual nutritional context. It moves video-derived behaviour from an endpoint of monitoring to a dynamic augmentation of physiology-based reference assessment.

The study makes four connected contributions. First, it defines an auditable seven-variable cow-day record joining video-derived feeding, rumination and lying durations with body weight, days in milk, parity and daily milk yield. Second, matched temporal and leave-one-cow-out comparisons isolate the incremental contribution of feeding duration to NRC-reference reconstruction. Third, animal-level error profiles delineate the supported operating domain and identify early lactation as a priority for targeted extension; two independent veterinary records add descriptive temporal context. Fourth, the cow-day indicator is placed within a staged digital-twin workflow linking implemented sensing and state construction to a specified operator-review gate.

Accordingly, the objectives were to (i) construct identity-associated cow-day records; (ii) evaluate behaviour-informed NRC-reference reconstruction under temporal and animal-held-out designs; (iii) isolate the incremental contribution of feeding duration after physiological inputs were represented; (iv) examine cow-level transfer and veterinary temporal context; and (v) map the analytical output to a staged operator-review workflow. Together, these objectives move dairy digital-twin research from recognising behaviour toward interpreting its day-to-day relationship with physiological context.

## 2. Study positioning

### 2.1 From behaviour recognition to farm decision support

Computer vision provides a passive and scalable means of observing multiple animals through shared infrastructure. YOLO-family detectors have been applied successfully to cattle localisation in complex barn environments, while tracking algorithms such as ByteTrack retain lower-confidence detections that might otherwise fragment trajectories during partial occlusion [15,16]. Video architectures including SlowFast and TimeSformer add the temporal information needed to distinguish behaviours expressed through sustained jaw movement, posture or activity sequences rather than through a single discriminating frame [17,18]. Together, these advances have made reliable behaviour recognition increasingly feasible under monitored farm conditions. The cow-day is the connecting unit that gives behaviour recognition agricultural meaning. Identity-associated events are aggregated over a biologically interpretable interval and aligned with the physiological and production history of the same animal. Data provenance, sensing-quality indicators and explicit handling of incomplete observations accompany that record, allowing the resulting signal to be traced from video event to nutritional context.

Alternative sensing modalities contribute complementary information. Accelerometers and commercial rumination collars can monitor feeding-related activity continuously and have been evaluated against direct observation, although their performance depends on device placement, behavioural definitions and farm conditions [13]. RFID-enabled feed bins and load-cell systems measure feeding visits and feed disappearance more directly and provide the preferred infrastructure for validating individual DMI models [12]. Video contributes a different set of advantages: non-contact deployment, simultaneous observation of multiple behaviours, visual context surrounding an event and the potential to use fixed barn-monitoring infrastructure. Its performance is shaped by illumination, occlusion, camera geometry and identity continuity. A robust smart-agriculture architecture should therefore integrate video as one component of a layered sensing system, alongside feed-weighing, wearable and production-record data where available.

### 2.2 Nutritional references, feeding behaviour and interpretation of agreement

Feeding time and DMI are biologically related but represent different quantities. Eating rate varies with body size, parity, ration composition, competition and meal structure [6,9–11,29]. Studies using measured intake have shown that feeding behaviour can improve intake models, although feeding duration alone cannot represent the full variation in consumption [29]. A single population-level relationship between time at the manger and DMI may therefore obscure meaningful differences among animals, particularly across body sizes and stages of lactation.

Physiological models approach the problem from a complementary direction. NRC (2001) estimates expected DMI from fat-corrected milk, metabolic body weight and an early-lactation adjustment [3]. Other nutritional systems integrate animal, ration and production characteristics to support precision feeding [20]. NASEM (2021) extends this framework by incorporating additional biological information, including parity-related and body-condition effects [21]. In the present dataset, body-condition score was recorded at two time points, whereas the analysis required a daily reference across the observation period. NRC (2001) was therefore selected as the temporally complete and reproducible reference compatible with the available longitudinal records.

NRC predictions describe population-based physiological expectations. Individual cows may depart from those expectations because of health, genotype, residual feed intake, ration properties, social conditions or management. Recursive structural models of residual feed intake address dependencies among intake, production, body-reserve change and efficiency at population or breeding scales [30]. These approaches are complementary to the present framework. They characterise longer-term biological covariance and genetic architecture, whereas the video layer contributes a continuously observed behavioural signal that can vary between updates of body weight or milk-composition records.

The analytical question is therefore whether video-derived feeding duration adds temporal information to physiology-based reference assessment. Because body weight, fat-corrected milk and days in milk are used to calculate the NRC reference, substantial agreement from models containing those variables is expected. The relevant evidence is the incremental change in held-out reference reconstruction when feeding duration is added under otherwise identical validation folds. Accordingly, lower prediction error is interpreted as closer reconstruction of the NRC reference rather than as demonstrated accuracy against consumed feed. Nested model comparisons and evaluation on entirely held-out animals establish the descriptive contribution and within-cohort transfer of the behavioural signal. Validation against individually measured intake represents the next biological evaluation stage.

### 2.3 Staged maturity of dairy digital-twin decision support

Digital twins originated in product-lifecycle management and industrial cyber-physical systems, where they describe synchronised computational representations used for monitoring, simulation and action [22–25]. Agricultural digital twins increasingly extend this principle to biological production systems, although their maturity depends on the evidence supporting each sensing, modelling, decision and action layer [31]. Animals add an important source of complexity because their responses to management depend on health, social context, ration palatability, previous exposure and physiological history. A credible livestock digital twin must therefore make the maturity and validation status of each component visible.

We organise the present system into four progressive levels: implemented sensing, implemented aggregation and state construction, prototype decision support and physical integration. At the first level, fixed video captured individual cow behaviour. At the second, identity-associated behavioural events were aggregated into daily cow records and aligned with physiological and production data. At the third, the NRC-referenced nutritional module generated cow-level deviation signals, while a previously described Unity prototype displayed cow avatars, current behaviour labels and a per-cow inspection panel through structured file exchange. An allocation rule was additionally examined through an offline mathematical plant to verify software behaviour under specified assumptions.

Together, these components establish the physical-to-digital state and decision-support stages of the dairy digital-twin pathway. Real-time application interfaces, operator-response logging, automated feeder integration and measured biological response constitute the subsequent stages toward bidirectional operation. This staged formulation allows the reader to distinguish farm-derived evidence from offline software evaluation while retaining a clear and testable route toward an operational digital twin.

### 2.4 Relationship to prior publications

The broader research programme has reported complementary elements of the dairy digital-twin pathway, and Table 1 distinguishes their respective contributions. Earlier work articulated conceptual architectures for livestock digital twins and precision dairy nutrition [26,27]. A separate article reported and evaluated the YOLOv11, ByteTrack and TimeSformer perception pipeline, structured event outputs, and a one-cow 24-hour illustration linking behaviour with intake-oriented inference and Unity visualisation [19]. A CCECE 2026 conference poster presented the same preliminary 24-hour architecture [28].

**Table 1.** Relationship of the present study to prior outputs from the research programme.

| Output | Scope already reported | Distinct scope of the present study |
| --- | --- | --- |
| Precision-nutrition digital-twin review [27] | Architectural taxonomy and implementation roadmap | Empirical cow-day analysis and evidence boundary |
| Video-perception article [19] | Perception evaluation, structured event records and one-cow 24-hour intake-oriented/Unity illustration | Sixteen-cow, 153-cow-day NRC-reference analysis, matched animal-held-out ablation and operating-domain evaluation |
| CCECE poster [28] | Conference-poster illustration of the preliminary 24-hour digital-twin architecture | Expanded NRC-reference analysis, 153 cow-days, operating-domain evaluation and maturity-accounted interpretation |

The present study begins after the perception stage and provides the substantive cow-level nutritional analysis. It extends the preliminary architecture through 153 cow-days, explicit daily state construction, temporal and leave-one-cow-out evaluation, matched physiology-plus-behaviour ablation, animal-level operating-domain analysis and a maturity-accounted decision-support workflow. The perception outputs are used as an established upstream data source, while the analytical contributions reported here concern their integration with physiological records and their interpretation within a nutritional-reference framework.

This evidence-aligned terminology unifies the research programme without conflating its stages: the present article contributes the 16-cow, 153-cow-day reference reconstruction and animal-transfer analysis, while the published perception layer remains its documented upstream source. The resulting framework is both extensible and auditable, two requirements for dairy digital twins intended to progress from continuous observation to responsible farm decision support.

## 3. Materials and methods

### 3.1 Study design, facility and ethical approval

The study was an exploratory, retrospective analysis of video-derived behaviour and routine farm records. Data originated from the Ruminant Animal Centre at Dalhousie University’s Agricultural Campus in Truro, Nova Scotia, Canada. The facility houses approximately 65 Holstein cows in a tie-stall barn. Each cow occupies an assigned stall with a lying area, a front feed manger and an adjacent water bucket. Cows received a total mixed ration comprising grass silage, corn silage, straw and concentrate, with composition adjusted through routine herd management. Milking occurred twice daily at approximately 04:30 and 16:00.

The barn used six Panasonic WV-S35302-F2L dome cameras and one Panasonic WV-X15700-V2L bullet camera. Cameras were mounted approximately 3-4 m above the barn, recorded continuously at 24 frames/s and provided infrared imagery during the five-hour dark period. The nutritional analysis used cows visible in three camera views. The tie-stall layout enabled stall-zone anchoring of identity, but it also constrained the study’s transferability to freestall environments.

All procedures complied with Canadian Council on Animal Care guidance and were approved by the Dalhousie University Animal Care and Use Committee (Protocol 2024-026; approved 16 May 2024). Video acquisition was passive and involved no additional animal contact.

### 3.2 Cohort and analytical records

Seventeen candidate lactating Holstein cows were video-aligned for up to 11 consecutive days in March 2025. Cow 403 was excluded before modelling because detection gaps produced atypically low apparent feeding time, leaving a 16-cow analytical cohort. The cohort spanned parities 1–7, body weights of 623–921 kg and 15–354 days in milk at the beginning of observation; mean daily milk yield was 43.6 kg/day. Five included cows were observed in Camera 001, six in Camera 002 and five in Camera 003. Cow-days coinciding with confirmed clinical events were separated from model fitting. The resulting primary dataset contained 153 clean cow-days; event-associated observations were retained only for the descriptive concordance analysis in Section 3.8.

For each cow-day, routine farm records supplied body weight, days in milk, parity, daily milk yield and milk fat percentage. Body weight was recorded during scheduled weighing and carried forward for no more than seven days. Days in milk increased deterministically by one per day; parity remained fixed during the study; morning and afternoon yields were summed. Milk fat percentage was used to calculate 4% fat-corrected milk. Table 2 summarises the cohort at the beginning of the observation window. Recording duration varied among cows; therefore, 16 cows multiplied by 11 days is a maximum rather than an initial complete-case denominator.

**Table 2.**
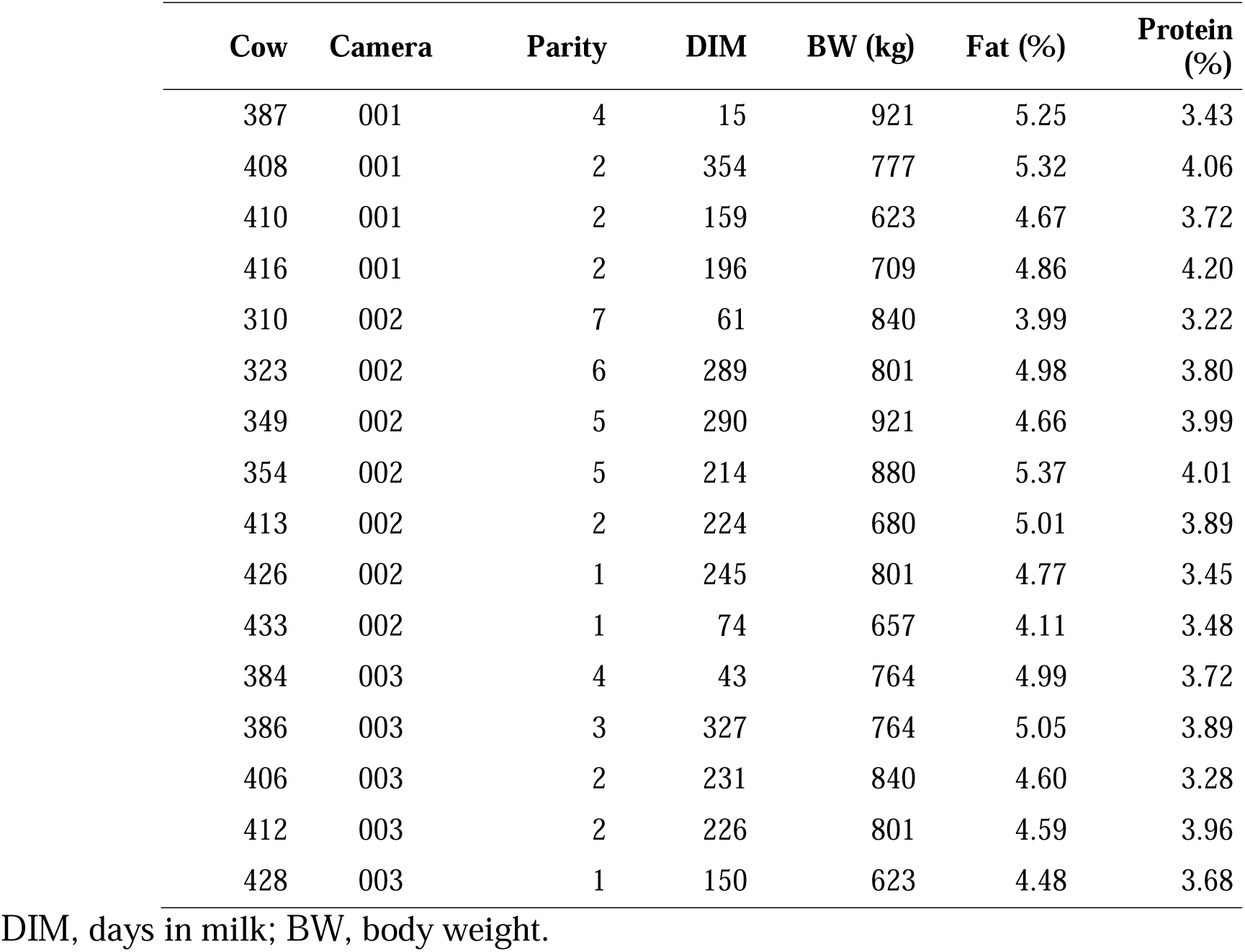
Physiological characteristics of the 16-cow nutritional cohort.

**Table 3.**
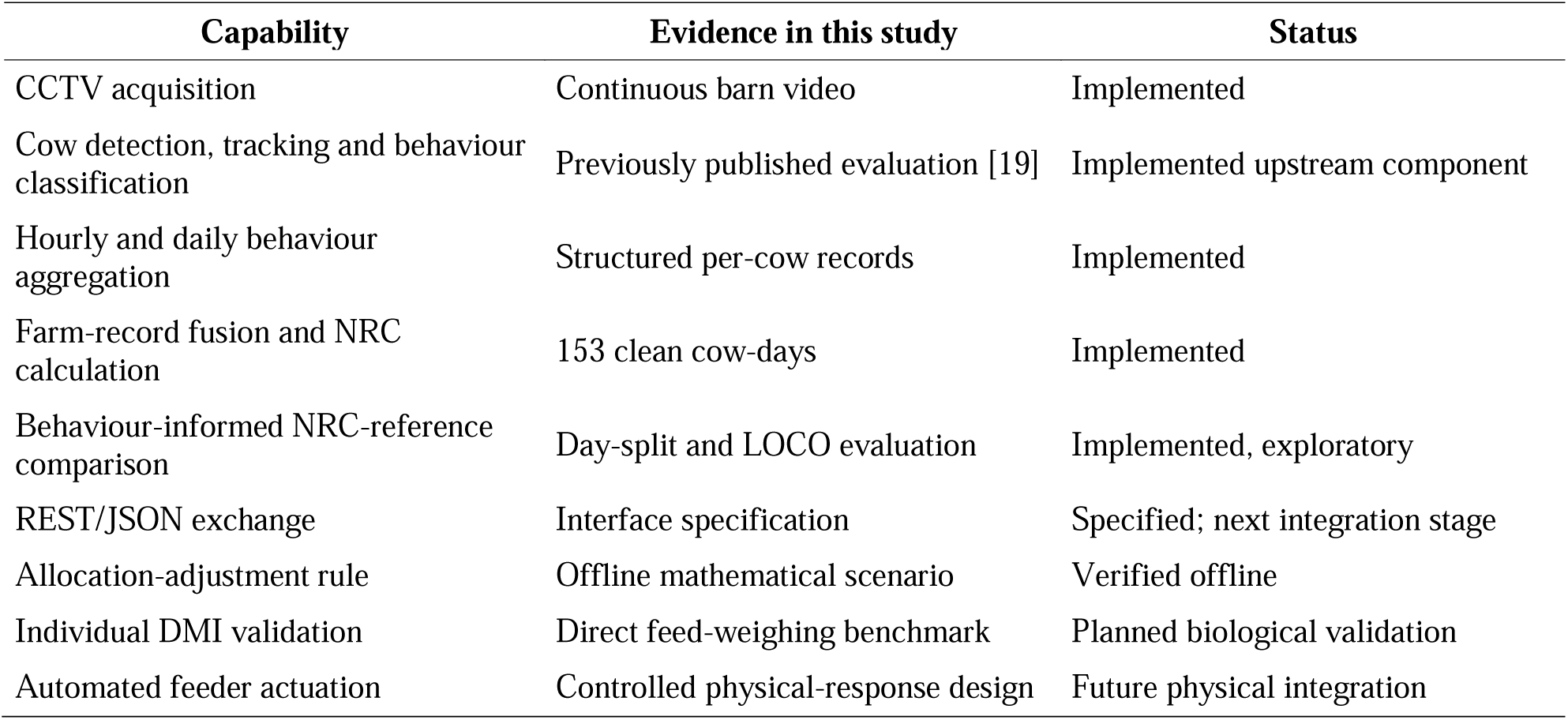
Evidence and implementation status of the proposed workflow.

| Capability | Evidence in this study | Status |
| --- | --- | --- |
| CCTV acquisition | Continuous barn video | Implemented |
| Cow detection, tracking and behaviour classification | Previously published evaluation [19] | Implemented upstream component |
| Hourly and daily behaviour aggregation | Structured per-cow records | Implemented |
| Farm-record fusion and NRC calculation | 153 clean cow-days | Implemented |
| Behaviour-informed NRC-reference comparison | Day-split and LOCO evaluation | Implemented, exploratory |
| REST/JSON exchange | Interface specification | Specified; next integration stage |
| Allocation-adjustment rule | Offline mathematical scenario | Verified offline |
| Individual DMI validation | Direct feed-weighing benchmark | Planned biological validation |
| Automated feeder actuation | Controlled physical-response design | Future physical integration |

### 3.3 Provenance of the video-derived behaviour records

The upstream perception pipeline has been published separately [19]. Briefly, YOLOv11 detected cows, ByteTrack associated detections through time, and stall-zone anchors maintained operational identity across file boundaries in the tie-stall layout. TimeSformer assigned one of seven clip labels: Standing, Lying, Drinking, Feeding and Standing, Feeding and Lying, Ruminating and Standing, or Ruminating and Lying. The published article documents the dataset, training design and perception performance; those established outputs provide the upstream input for the present cow-day analysis [19].

Each accepted clip produced an event record of the form {cow identifier, camera identifier, behaviour label, start time, end time, duration, confidence}. Predictions with confidence below 0.60 were excluded before aggregation. A three-clip majority rule reduced isolated label changes. Feeding duration combined Feeding and Standing with Feeding and Lying; rumination combined both rumination postures; lying duration combined all labels that included recumbency. Because these categories can overlap biologically, a ruminating-and-lying interval contributed to both the rumination and lying totals.

Stable stall zones supplied the operational identity anchor for the present tie-stall records. Transfer to group-housed barns is a distinct engineering stage requiring full-day IDF1 and HOTA reporting, cross-camera evaluation and re-identification based on appearance, geometry or sparse RFID confirmation [32–34]. Propagating identity confidence into the cow-day record will preserve traceability between behaviour and the correct physiological profile.

### 3.4 Daily cow representation and data provenance

Clip-level events were summed to hourly intermediates and then to daily totals. The seven-field cow representation at day *t* was

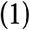

where the first three terms are video-derived durations in minutes/day; BW is body weight in kg; DIM is days in milk; parity is the lactation number; and daily milk yield is expressed in kg/day. Milk fat percentage was retained as an auxiliary farm-record field used to derive fat-corrected milk. Feeding duration served as the behaviour term in the nutritional model, while rumination and lying remained contextual state variables that preserve the framework’s capacity for future multivariate evaluation.

The prototype exchanged structured cow-event records through CSV files and retained predictions at or above 0.60 confidence. These implementation choices supported reproducible aggregation during the study. The next deployment iteration adds a load-tested real-time interface, an explicit unobserved state and a valid-video coverage field so that low-coverage cow-days can be routed directly to review.

### 3.5 NRC reference calculation

The NRC (2001) reference was calculated from routine physiological records [3]. Four-percent fat-corrected milk was

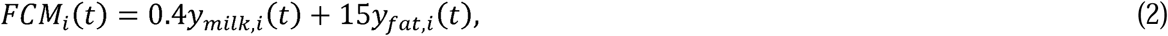

With

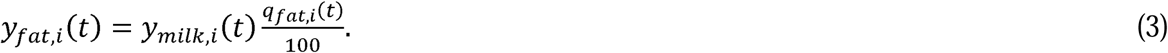

The corresponding reference DMI was

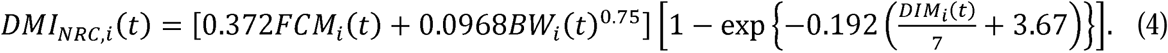

Equation (4) serves as the NRC-derived physiological reference compatible with the daily farm records. The reported RMSE, MAE, bias and MAPE therefore quantify reference reconstruction; direct individual feed weighing is the prospective biological validation standard.

### 3.6 Behaviour-informed models

Four comparisons were used to separate model form from behavioural contribution.

#### 3.6.1 Feeding-duration baseline

The simplest model assumed a constant eating rate,

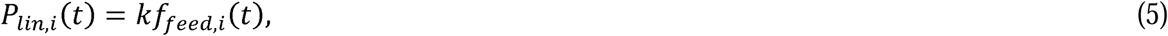

where *P_lin_* is expressed on the kg DM/day scale of the NRC reference. The coefficient *k* was refitted within each training fold. This model provides a deliberately simple benchmark, not a mechanistic representation of intake.

#### 3.6.2 Reduced behaviour-informed index

A secondary reduced index combined the daily behavioural signal with body size and production information:

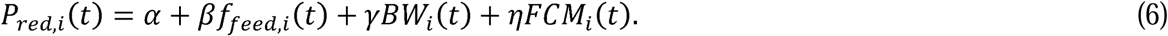

When fitted on all 153 clean cow-days for descriptive calibration, Eq. (6) was

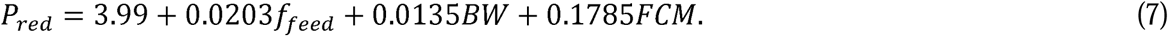

Equation (7) is an NRC-aligned index in which physiology establishes the broad expected level and daily feeding duration provides the time-varying behavioural term. Its animal-transfer performance was evaluated through leave-one-cow-out refitting, described in Section 3.7.

#### 3.6.3 Primary matched behavioural-augmentation analysis

To ask whether feeding duration added information beyond the variables that construct the NRC reference, four ordinary least-squares variants were compared under the same leave-one-cow-out partitions:

1. an intercept-only population-mean model;
2. a physiology-only model using *BW, FCM* and *DIM*;
3. a behaviour-only model using *f_feed_*; and
4. a combined model using *BW, FCM, DIM* and *f_feed_*.

The combined model is the primary matched behavioural-augmentation analysis; Eq. (6) is retained as a secondary reduced index. Because BW, FCM and DIM define the NRC target, strong performance of the physiology-only and combined models is expected. Indeed, applying Eq. (4) directly would reproduce the target exactly by definition. The relevant result is the incremental held-out change when feeding duration is added under otherwise identical folds. No inferential p-value is used to imply independence among repeated cow-days.

#### 3.6.4 Saturating sensitivity model

A Michaelis-Menten form was evaluated as a secondary biological sensitivity analysis:

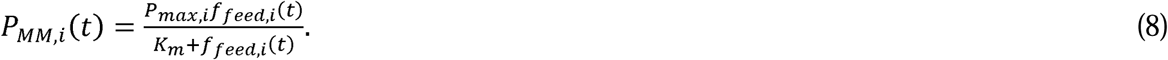

The half-saturation value *K_m_* was searched over 10-300 min/day, with 30-150 min/day treated as biologically plausible. The selected value was 30 min/day. The cow-specific ceiling *P_max,i_* was derived from each animal’s mean NRC reference and mean feeding duration. This construction anchors the curve to the cow’s own history and therefore favours agreement at the mean. It also prevents honest leave-one-cow-out application to a completely new animal without a calibration period. The model was consequently retained for day-split sensitivity and the offline plant illustration, not presented as the principal deployable result.

### 3.7 Held-out evaluation and statistical reporting

Two held-out strategies addressed different sources of leakage. In the temporal day-split, each cow’s clean days were ordered and divided approximately in half. Coefficients were fitted to the earlier days pooled across cows and evaluated on the later days. This tests transfer across time but allows each test cow to contribute earlier observations to training.

Leave-one-cow-out cross-validation (LOCO) provided the primary evaluation for the matched ablation and the strict transfer evaluation for the reduced index. In each of 16 folds, all cow-days from one cow were withheld, the model was fitted on the other 15 animals and predictions were generated for every day of the held-out cow. Pooled LOCO RMSE, MAE, bias, MAPE and R² were calculated from the combined held-out predictions. Per-cow errors are also reported so that dependence among repeated days and heterogeneity across animals remain visible. LOCO prevents an animal’s data from entering the coefficients used to predict that animal, but it does not make observations within a cow independent. Inferences beyond this herd should therefore be based on the 16 animal-level folds rather than on 153 nominally independent rows.

Metrics were defined as

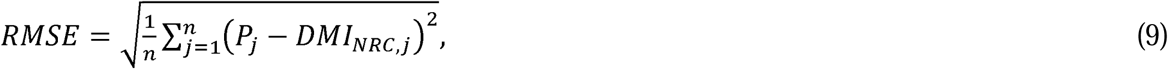

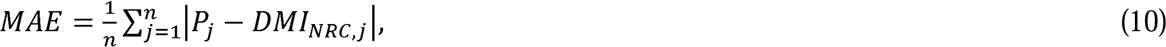

And

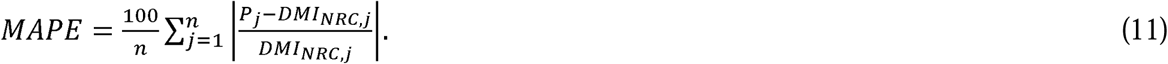

Bias was the mean signed difference between the model prediction and the NRC reference. The coefficient of determination was calculated against the held-out NRC values. For the reduced-index LOCO analysis, heterogeneity is reported through all 16 held-out-cow errors. The archived matched ablation supports pooled held-out metrics only; a cow-clustered interval was not available.

### 3.8 Descriptive comparison with veterinary records

Two veterinary events occurred during the wider observation window: a recurrence of a teat injury in Cow 354, recorded on 20 March 2025 and treated with Anafen, and an event recorded as hypocalcaemia (“milk fever”) in Cow 349 on 23 March 2025, treated with intravenous calcium and Meloxidyl. Event identity and date were not used as model inputs. Feeding duration and the ratio *P_MM_/DMI_NRC_* were plotted around each event to determine whether an unusual trajectory was temporally concordant with the independently entered record.

This post hoc descriptive comparison examined temporal concordance between independently entered veterinary records and unusual cow-level trajectories. The two cases define a prospective evaluation question for sensitivity, specificity, lead time and false-alarm burden rather than a diagnostic endpoint.

### 3.9 Software and reproducibility

Model fitting and evaluation used Python, NumPy, pandas and scikit-learn. The published perception pipeline used PyTorch, Ultralytics YOLO, ByteTrack and OpenCV [19]. The behaviour-visualisation prototype used Unity 2022 LTS. Code for the published upstream perception pipeline is available at https://github.com/mooanalytica/digital-twin-dairycow. Availability of the daily analytical records and downstream scripts is stated in the Data availability section.

## 4. Results

### 4.1 Composition and reference range

The 153 clean cow-days represented a heterogeneous single-herd cohort. Cows ranged from 623 to 921 kg and from 15 to 354 days in milk. The resulting NRC reference values spanned approximately 17.9-37.3 kg DM/day, while observed feeding durations spanned approximately 76-442 min/day for most clean observations. One excluded animal had severe video-coverage gaps. This exclusion illustrates why an unobserved state and coverage-aware output withholding are necessary in any deployment: zero duration cannot safely be treated as zero behaviour when the camera record is incomplete.

### 4.2 Primary analysis: behavioural augmentation of physiological expectation

The primary analysis asked whether video-derived feeding duration added held-out information after the physiological variables used to construct the NRC reference were represented. Four ordinary least-squares variants were compared under identical leave-one-cow-out (LOCO) partitions. In every fold, all records from one cow were withheld, coefficients were fitted on the remaining 15 cows and predictions were generated for every cow-day of the held-out animal.

The population-mean model achieved RMSE 3.009 kg DM/day, MAE 2.359 kg DM/day, MAPE 8.5% and R² = −0.116. Feeding duration alone performed similarly, with RMSE 2.913 kg DM/day, MAE 2.250 kg DM/day, MAPE 8.1% and R² = −0.045, showing that time at the manger cannot reconstruct physiological expectation by itself. The physiology-only model, using body weight, fat-corrected milk and days in milk, achieved RMSE 2.015 kg DM/day, MAE 1.267 kg DM/day, MAPE 4.5% and R² = 0.500. Adding video-derived feeding duration reduced RMSE to 1.763 kg DM/day, MAE to 1.159 kg DM/day and MAPE to 4.2%, while R² increased to 0.617 (Table 4; Figure 2).

**Figure 1.**
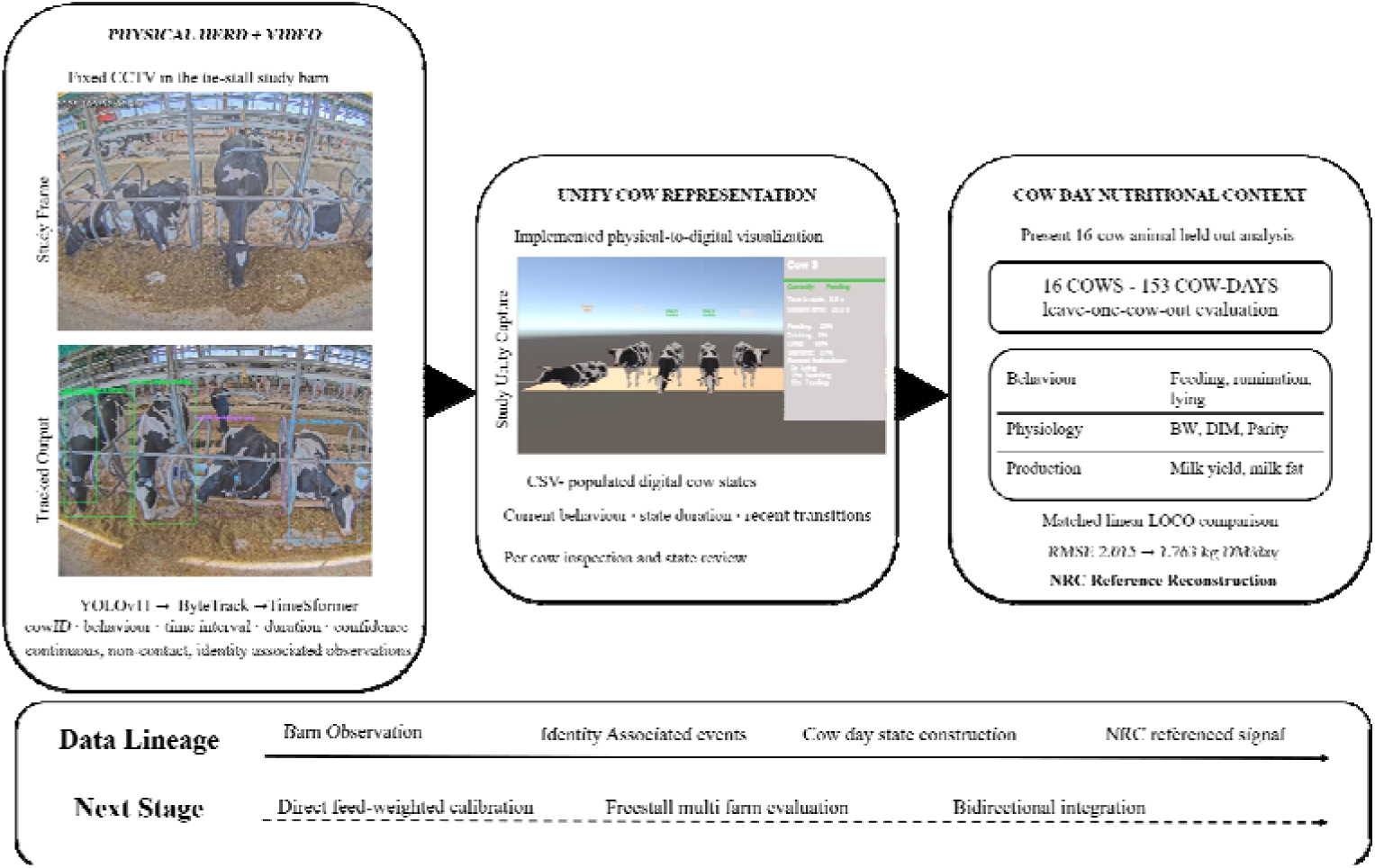
From the physical herd to cow-level digital-twin context. Existing barn video supplies continuous, non-contact observation. The evaluated YOLOv11-ByteTrack-TimeSformer layer [19] associates events with individual cows; this study fuses those records with body weight, days in milk, parity, milk yield and milk fat to construct auditable cow-day states. Across 16 cows and 153 cow-days, the digital counterparts support animal-held-out NRC-reference analysis; the resulting signals are designed for focused operator review. Solid elements trace the evaluated analytical pathway; the dashed pathway shows translation toward operator use, measured intake, prospective farm evaluation and bidirectional operation.

**Figure 2.**
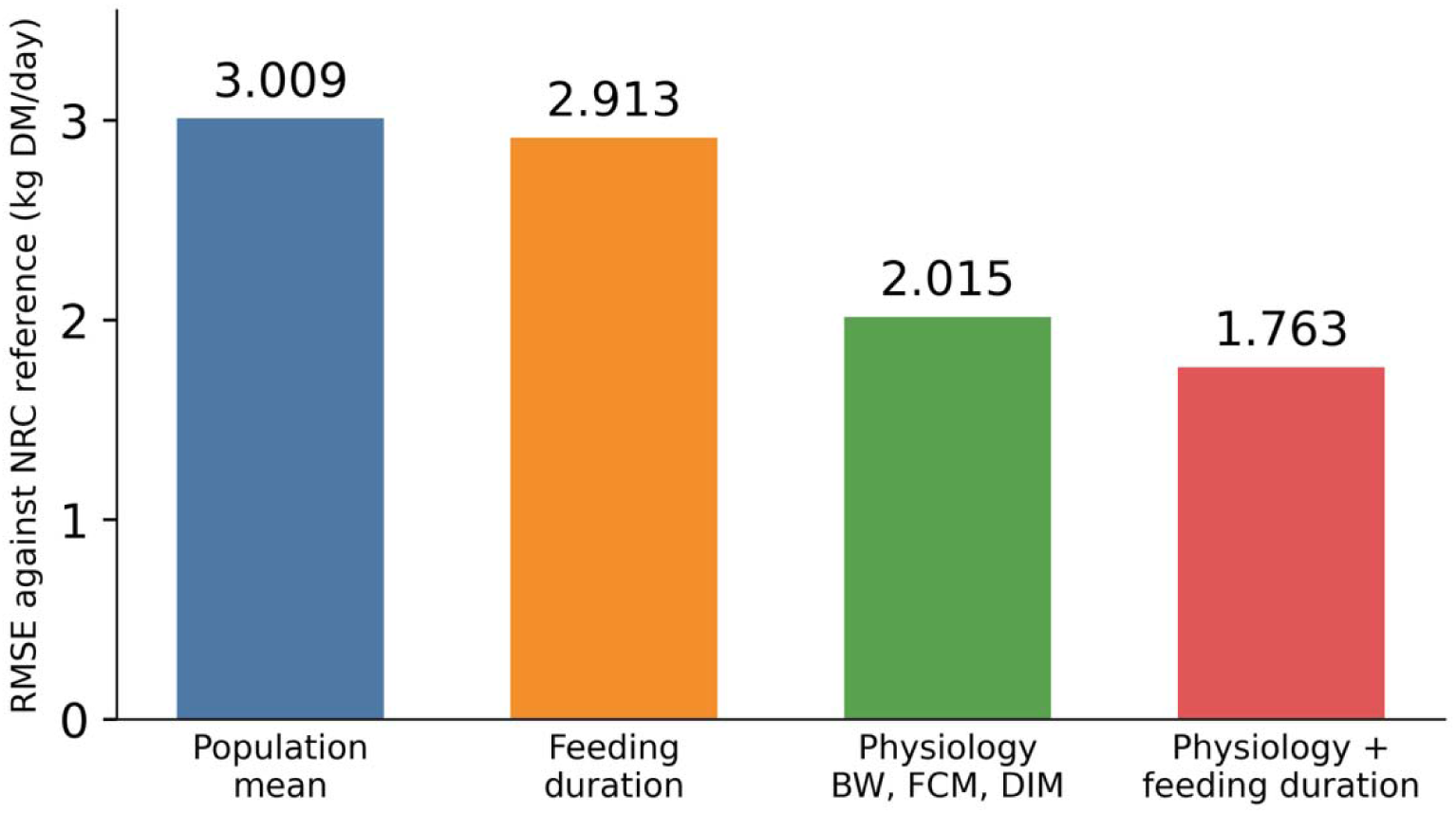
Matched animal-held-out ablation of NRC-reference reconstruction (153 cow-days; 16 folds). The physiology models used identical LOCO splits and feeding duration was the sole added predictor. Values are pooled descriptive errors from held-out cow-days against the model-derived NRC reference.

**Table 4.** Primary matched leave-one-cow-out ablation against the NRC (2001) reference (153 cow-days)

| Predictor set | RMSE (kg/day) | MAE (kg/day) | MAPE (%) | $R^2$ |
| --- | --- | --- | --- | --- |
| Population mean | 3.009 | 2.359 | 8.5 | -0.116 |
| Feeding duration only | 2.913 | 2.250 | 8.1 | -0.045 |
| Physiology only: BW, FCM, DIM | 2.015 | 1.267 | 4.5 | 0.500 |
| Physiology plus feeding duration | 1.763 | 1.159 | 4.2 | 0.617 |

The matched addition of feeding duration corresponded to reductions of 0.252 kg DM/day in pooled held-out RMSE, 0.108 kg DM/day in MAE and 0.3 percentage points in MAPE. The physiological variables account for most NRC-reference alignment, as expected from the construction of the reference. The separately sensed feeding term supplies the day-varying component. In the pooled descriptive fit, its coefficient was +0.0159 kg DM per minute. Cow-clustered inferential uncertainty was unavailable, so this coefficient is reported descriptively and no population-level significance is claimed.

### 4.3 Secondary reduced index and temporal transfer

A reduced behaviour-informed index using feeding duration, body weight and fat-corrected milk was retained as a secondary analysis. Under a temporal split in which earlier observations were used for fitting and later observations for testing, this index achieved RMSE 1.811 kg DM/day, MAE 1.377 kg DM/day, bias −0.196 kg DM/day, MAPE 4.9% and R² = 0.663. Under the stricter LOCO design, pooled RMSE was 2.362 kg DM/day, MAE 1.605 kg DM/day, bias +0.229 kg DM/day, MAPE 5.8% and R² = 0.313 (Table 5; Figure 3).

**Figure 3.**
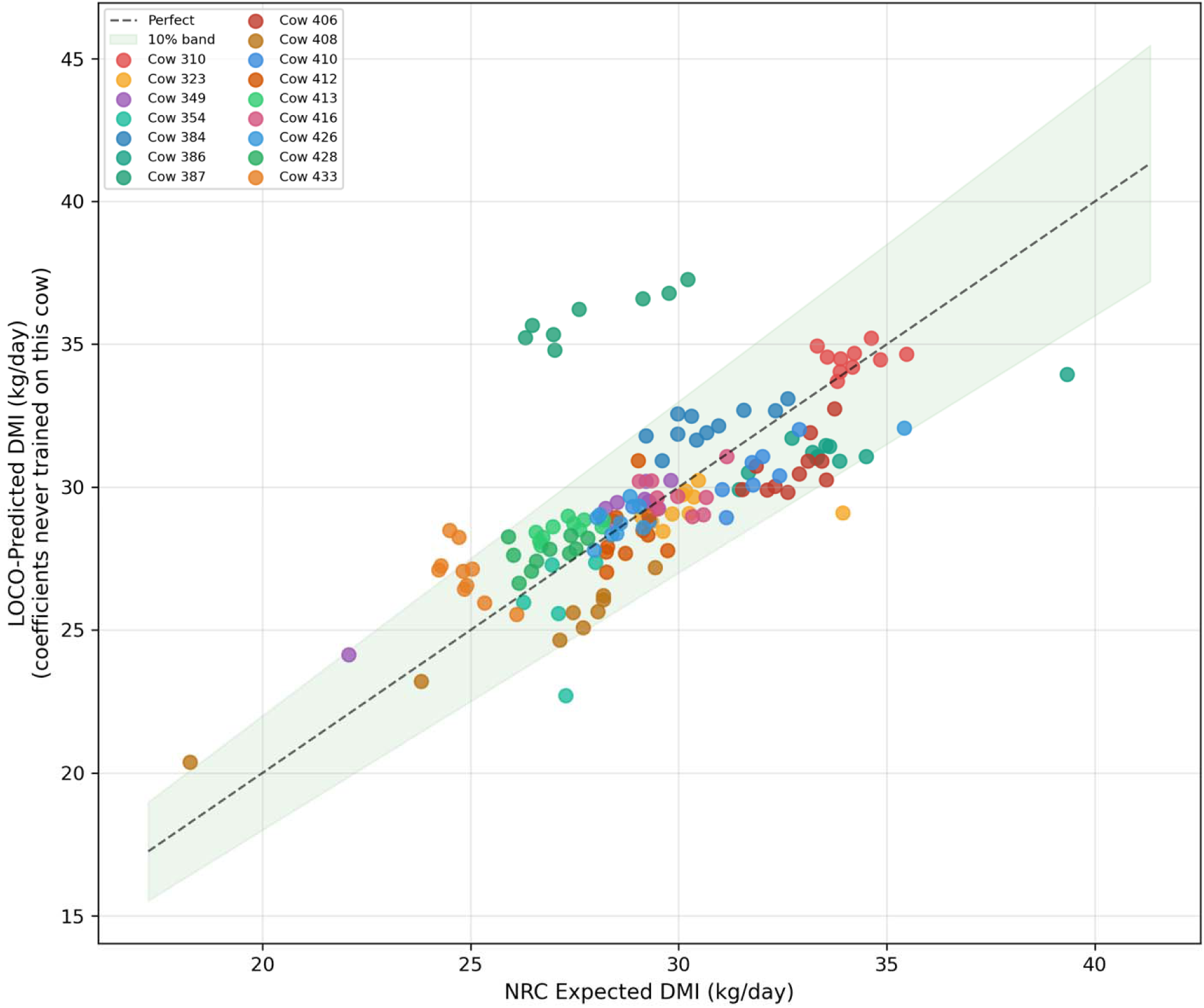
Animal-held-out predictions from the reduced behaviour-informed index versus the NRC (2001) reference (16 cows; 153 cow-days). Each prediction was generated without the corresponding cow in model fitting. Cow 387 entered at 15 days in milk, revealing the fresh-cow operating-domain boundary; the identity line and ±10% band describe NRC-reference agreement.

**Table 5.** Secondary held-out analyses against the NRC (2001) reference.

| Model | Evaluation | n cow-days | RMSE (kg/day) | MAPE (%) |  |
| --- | --- | --- | --- | --- | --- |
| Feeding baseline | Temporal test | 81 | 5.773 | 16.1 | -2.428 |
| Feeding baseline | LOCO | 153 | 5.539 | 15.8 | -2.779 |
| Reduced model, Eq. (6) | Temporal test | 81 | 1.811 | 4.9 | 0.663 |
| Reduced model, Eq. (6) | LOCO | 153 | 2.362 | 5.8 | 0.313 |
| Saturating sensitivity model | Temporal test | 81 | 1.578 | 3.7 | 0.744 |

**Table 6.** Per-cow LOCO performance of the secondary reduced behaviour-informed index.

| **Cow** | **Camera** | **RMSE (kg/day)** | **MAPE (%)** |
| --- | --- | --- | --- |
| 310 | 002 | 0.681 | 1.6 |
| 323 | 002 | 1.584 | 3.2 |
| 349 | 002 | 1.069 | 3.5 |
| 354 | 002 | 2.059 | 5.3 |
| 384 | 003 | 1.530 | 4.7 |
| 386 | 003 | 2.481 | 6.8 |
| 387 | 001 | 7.770 | 28.8 |
| 406 | 003 | 2.254 | 6.8 |
| 408 | 001 | 2.000 | 7.6 |
| 410 | 001 | 1.280 | 3.8 |
| 412 | 003 | 0.989 | 2.9 |
| 413 | 002 | 1.223 | 4.5 |
| 416 | 001 | 0.829 | 2.4 |
| 426 | 002 | 1.150 | 2.3 |
| 428 | 003 | 0.992 | 3.2 |
| 433 | 002 | 2.322 | 8.8 |

The feeding-duration baseline showed substantially weaker transfer under the same partitions: LOCO RMSE 5.539 kg DM/day, MAPE 15.8% and R² = −2.779. The pattern was consistent with a single eating-rate coefficient failing to represent cows across an approximately 300 kg body-weight range. A cow-calibrated saturating model achieved temporal-test RMSE 1.578 kg DM/day, MAPE 3.7% and R² = 0.744, but its cow-specific ceiling required within-animal historical calibration. It was therefore treated as a sensitivity analysis rather than a model for previously unseen cows.

### 4.4 Cow-level variation and operating domain

The reduced index’s pooled LOCO performance concealed substantial animal-level variation. Median per-cow RMSE was 1.405 kg DM/day (interquartile range 1.050-2.108), and median per-cow MAPE was 4.15% (interquartile range 3.13-6.80). Cow 387, observed from 15 days in milk and representing the least densely sampled lactation region, had RMSE 7.770 kg DM/day and MAPE 28.8% (Figure 4).

**Figure 4.**
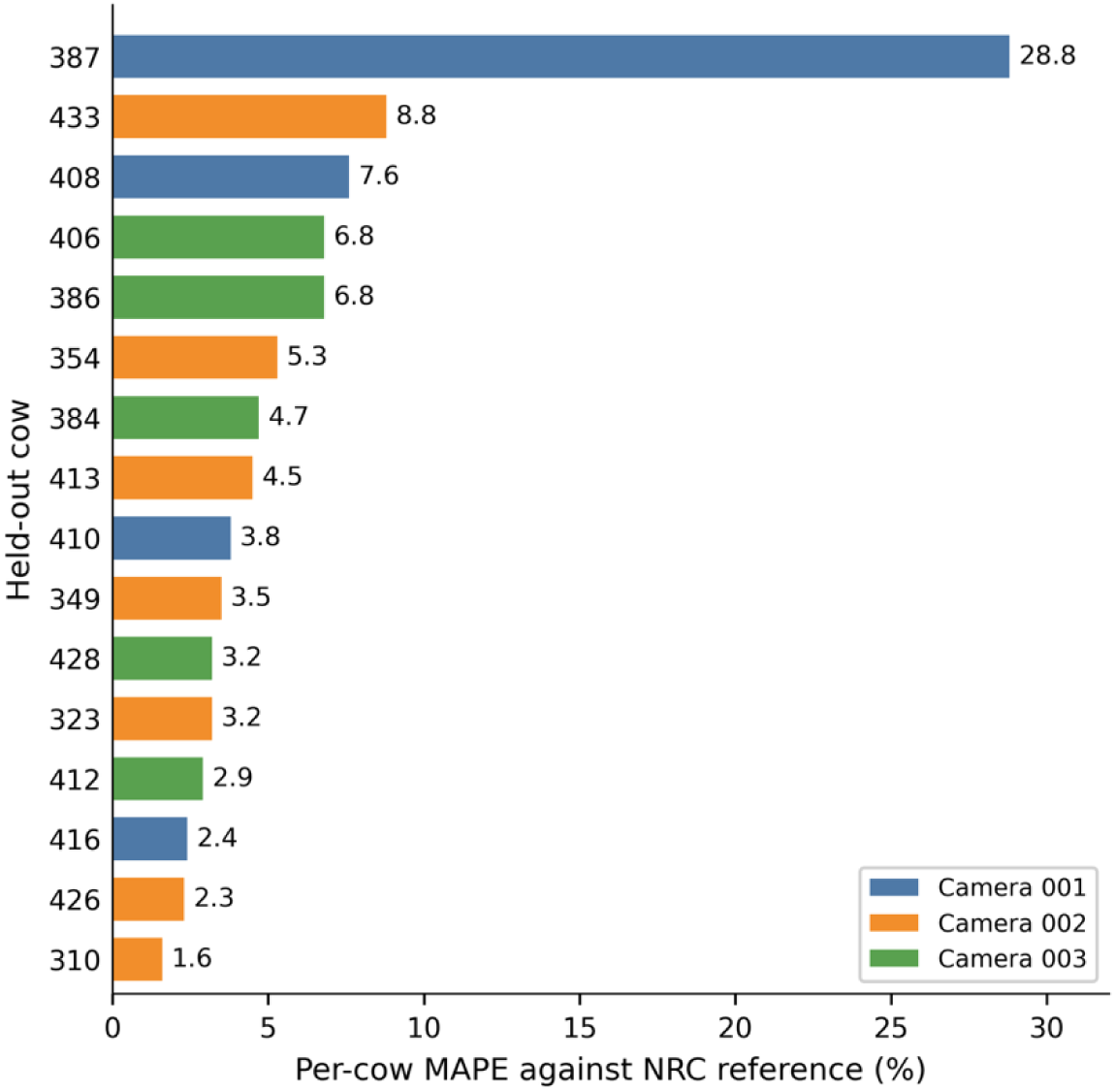
Per-cow held-out MAPE for the reduced body-weight, fat-corrected-milk and feeding-duration index. The plot makes transfer heterogeneity visible and identifies Cow 387, the only animal entering at 15 days in milk, as the fresh-cow operating-domain extension.

When Cow 387 was withheld, the feeding-duration coefficient fell to 0.0047 kg/min, approximately one quarter of the 0.020-0.023 kg/min range observed in typical folds, while the FCM coefficient increased to 0.271. The fold therefore exposed sensitivity to training composition rather than a generic early-lactation effect. This fresh-cow case lies outside the model’s currently supported operating domain and motivates stage-specific calibration or explicit abstention when days in milk and related variables fall outside the densely represented training range.

### 4.5 Descriptive concordance with independent veterinary records

Two independently recorded veterinary events coincided with unusual feeding-derived trajectories. Cow 354’s feeding duration fell to 75.6 min on 19 March, its lowest observed daily value, one day before treatment for a recurrent teat injury on 20 March. The corresponding cow-calibrated sensitivity-model/reference ratio was 0.81. Cow 349 showed a ratio of 1.21 on 22 March, one day before treatment for a farm record entered as hypocalcaemia on 23 March. Health status and treatment date were absent from model inputs. The deviations occurred in opposite directions, favouring interpretation as departures from recent cow-level patterns rather than a universal disease threshold. With two events, the analysis provides descriptive biological plausibility and motivates prospective trajectory-based evaluation; it does not estimate diagnostic sensitivity, specificity or false-alarm burden.

## 5. Discussion

### 5.1 Behaviour as a dynamic layer around physiological expectation

This study addresses a persistent gap between animal monitoring and biologically interpretable decision support. Computer vision can generate continuous, identity-associated activity records, while nutritional equations describe physiological expectation. By combining them at the cow-day level, the framework turns feeding behaviour from a classification endpoint into a dynamic signal around physiological context.

The matched LOCO ablation provides the clearest evidence for that contribution. Body weight, fat-corrected milk and days in milk encoded much of the NRC reference by construction, producing RMSE 2.015 kg DM/day. Adding feeding duration reduced RMSE to 1.763 kg DM/day and increased R² from 0.500 to 0.617. The magnitude is moderate; its importance lies in supplying an independently sensed variable that can change daily while body weight and some production records remain stable. Video therefore adds temporal responsiveness to a predominantly physiological reference.

The appropriate interpretation is an NRC-referenced behavioural deviation signal. Feeding duration captures a day-varying component of the cow’s interaction with feed, while the physiological reference describes expected demand. The signal can prioritize cow-days for contextual review alongside ration delivery, refusals, milk records, health history and sensing quality. Direct feed weighing will determine how closely this behavioural augmentation tracks consumption.

The two veterinary cases illustrate how that review could be contextualized. Both showed conspicuous departures before independently entered treatment dates, and the departures occurred in opposite directions. An informative alert should therefore communicate the direction, magnitude and persistence of change together with recent baseline, video coverage and model confidence. As a descriptive two-event comparison, these cases nominate a prospective evaluation question.

### 5.2 Agricultural significance and complementary sensing

Video contributes a distinctive combination of non-contact measurement, shared infrastructure and behavioural context. It can distinguish posture and position relative to the manger, preserve an auditable visual record and observe several cows through one camera network. These properties make it attractive where wearable deployment is burdensome or fixed barn-monitoring infrastructure is available. The cow-day architecture also demonstrates how continuous behaviour can be integrated with routine farm records without collapsing either source into an opaque score.

Its greatest value will emerge in combination with other technologies. Instrumented feed bins and machine-vision measurements of feed disappearance remain essential for direct DMI calibration [12,35]. Accelerometers and rumination collars provide complementary evidence during darkness or visual occlusion [13]. Agreement across video, wearables and feed measurements can strengthen confidence; disagreement can identify sensing faults or unusual biological states. A digital twin should preserve that provenance so an operator can distinguish whether a deviation originates in observed behaviour, a physiological record or the sensing layer. Rumination and lying were retained in the daily cow record but excluded from the nutritional regression. This choice favoured parsimony and avoided introducing poorly identified associations into 153 repeated observations. Both behaviours remain valuable for health and welfare monitoring and can support future multivariate deviation models when larger, prospectively labelled datasets become available. Their inclusion in the state representation supports extensibility without overstating their present nutritional role.

### 5.3 Operating domain, identity and transfer

Animal-held-out evaluation delineated the current operating domain as well as pooled performance. Cow 387, the only animal entering at 15 days in milk, had MAPE 28.8%, whereas the remaining cohort occupied a better-supported physiological region. This observation identifies early lactation as a priority for targeted data collection and stage-specific evaluation. Distribution-aware output gating can route sparsely represented cow states to review until that extension is available. Stable tie-stall locations supplied a practical identity anchor for the present cow-day records. Extending the architecture to freestall systems will require full-day IDF1 and HOTA reporting, cross-camera evaluation and re-identification based on appearance, geometry or sparse RFID confirmation [32–34]. Propagating identity confidence into the cow-day record will preserve the framework’s central strength: traceability between behavioural history and the correct physiological profile.

LOCO demonstrated transfer between cows within one herd while preventing a held-out animal from contributing to its fitted coefficients. A logical next external evaluation is a multi-farm design spanning housing, ration, season, lactation stage and camera geometry. Reporting both cow-level and farm-level performance will distinguish genuine transfer from increases in correlated cow-days.

### 5.4 A staged pathway toward an operational dairy digital twin

The implemented system occupies a useful intermediate stage in dairy digital-twin development. Continuous sensing populates an explicit cow-level representation; farm records supply physiological context; an NRC-referenced analysis generates a deviation signal; and the architecture specifies a gate for operator review and abstention. These functions advance beyond an isolated behaviour classifier while preserving a clear distinction between state interpretation and physical intervention. Framing the system as a staged dairy digital twin makes each layer independently testable and prevents nominal feedback arrows from substituting for biological evidence.

Three steps would strengthen the pathway to operation. First, individual feed disappearance should be measured so calibration and uncertainty can be evaluated against observed DMI. Second, the full perception-to-record pathway should be tested prospectively in freestall and multi-farm settings, with identity and missing-data uncertainty propagated into daily outputs. Third, any allocation guidance should be compared with nutritionist-led management in a preregistered, welfare-monitored design incorporating safety limits, operator override and explicit withholding rules.

The broader principle extends beyond dairy nutrition. Smart-agriculture systems commonly combine high-frequency sensors with lower-frequency biological or management references. The key question is not only whether a sensor predicts a target, but whether it contributes new temporal information, exposes its provenance and abstains beyond the supported domain. This framework provides a practical template for that form of evidence-calibrated integration.

## 6. Conclusions

By transforming identity-associated video events into physiologically contextualized cow-day states, this study advances dairy video analytics from behaviour recognition towards biological interpretation. Each physical cow was represented at a daily cadence by a traceable computational state combining feeding, rumination and lying behaviour with body weight, lactation stage, parity and milk production. This state-and-interpretation layer establishes an essential physical-to-digital foundation for a dairy digital twin and enables analysis at the level most relevant to farm management: the individual cow.

Across 16 Holstein cows and 153 cow-days, adding video-derived feeding duration to a matched linear model containing body weight, fat-corrected milk and days in milk reduced animal-held-out RMSE against the NRC reference from 2.015 to 1.763 kg DM/day, a 12.5% reduction. MAE decreased from 1.267 to 1.159 kg DM/day, MAPE declined from 4.5% to 4.2%, and R² increased from 0.500 to 0.617. Because identical physiological predictors and leave-one-cow-out folds were used, this comparison isolates the contribution of the behavioural term within the evaluated linear framework. These metrics quantify augmentation of an NRC-derived reference rather than accuracy against observed individual intake. The cow-level analysis was equally informative: the sole animal entering the study at 15 days in milk showed elevated error, identifying this underrepresented fresh-cow case as a priority for stage-aware calibration and evidence-based output gating.

The central contribution is therefore the bridge between sensing and biological context, rather than either component in isolation. Continuous observations are converted into identity-linked records, anchored to physiological expectation and carried with their provenance towards a specified operator-review gate. This auditable pathway provides a practical foundation for direct-intake calibration, freestall identity evaluation, multi-farm transfer testing and prospective bidirectional integration.

More broadly, the study offers an evidence-based blueprint for dairy digital twins: their agricultural value emerges when a digital counterpart does more than mirror an animal, instead organising continuous observations into biologically interpretable, cow-specific evidence for timely and accountable decisions.

## Declarations Ethics approval

All animal-management procedures complied with Canadian Council on Animal Care guidelines and were approved by the Dalhousie University Animal Care and Use Committee (Protocol 2024-026; approval date 16 May 2024). Data acquisition was passive and non-invasive.

## CRediT authorship contribution statement

**Shreya Rao:** Methodology, Software, Validation, Formal analysis, Investigation, Data curation, Visualization, Writing - original draft. **Suresh Raja Neethirajan:** Conceptualization, Methodology, Supervision, Resources, Funding acquisition, Project administration, Writing - review and editing.

## Data availability

Code for the previously published perception pipeline is available at https://github.com/mooanalytica/digital-twin-dairycow under the MIT licence. The cow-level farm and veterinary records contain institutionally governed animal-research data. A de-identified analytical table and the nutritional-modelling and simulation scripts are available from the corresponding author on reasonable request, subject to institutional approval and applicable data-governance requirements.

## Acknowledgements

The authors thank the staff and animal caretakers of the Ruminant Animal Centre, Dalhousie University, for assistance with data collection and routine herd records. The authors also acknowledge Eduardo Garcia for contributions to the previously published perception pipeline cited in this work.

## References

[1] J. Lindblom, C. Lundström, M. Ljung, A. Jonsson, Promoting sustainable intensification in precision agriculture: review of decision support systems development and strategies, Precision Agriculture 18 (2017) 309–331. 10.1007/s11119-016-9491-4

[2] S. Neethirajan, B. Kemp, Digital Livestock Farming, Sensing and Bio-Sensing Research 32 (2021) 100408. 10.1016/j.sbsr.2021.100408.

[3] National Research Council, Nutrient Requirements of Dairy Cattle, 7th rev. ed., National Academies Press, Washington, DC, 2001. 10.17226/9825.

[4] K.A. Beauchemin, Invited review: Current perspectives on eating and rumination activity in dairy cows, Journal of Dairy Science 101 (2018) 4762–4784. 10.3168/jds.2017-13706.

[5] C. Johnston, T.J. DeVries, Short communication: Associations of feeding behavior and milk production in dairy cows, Journal of Dairy Science 101 (2018) 3367–3373. 10.3168/jds.2017-13743.

[6] M.S. Allen, Drives and limits to feed intake in ruminants, Animal Production Science 54 (2014) 1513–1524. 10.1071/AN14478.

[7] J.K. Drackley, Biology of Dairy Cows During the Transition Period: the Final Frontier?, Journal of Dairy Science 82 (1999) 2259–2273. 10.3168/jds.S0022-0302(99)75474-3.

[8] R. Antanaitis, K. Džermeikaitė, J. Krištolaitytė, I. Ribelytė, A. Bespalovaitė, D. Bulvičiūtė, A. Rutkauskas, Alterations in Rumination, Eating, Drinking and Locomotion Behavior in Dairy Cows Affected by Subclinical Ketosis and Subclinical Acidosis, Animals 14 (2024) 384. 10.3390/ani14030384.

[9] T.J. DeVries, M.A.G. Von Keyserlingk, D.M. Weary, K.A. Beauchemin, Measuring the Feeding Behavior of Lactating Dairy Cows in Early to Peak Lactation, Journal of Dairy Science 86 (2003) 3354–3361. 10.3168/jds.S0022-0302(03)73938-1.

10. M.A.G. Von Keyserlingk, D.M. Weary, Review: Feeding behaviour of dairy cattle: Measures and applications, Can. J. Anim. Sci. 90 (2010) 303–309. 10.4141/CJAS09127.

[11] R.J. Grant, J.L. Albright, Effect of Animal Grouping on Feeding Behavior and Intake of Dairy Cattle, Journal of Dairy Science 84 (2001) E156–E163. 10.3168/jds.S0022-0302(01)70210-X.

[12] I. Halachmi, Y. Edan, U. Moallem, E. Maltz, Predicting Feed Intake of the Individual Dairy Cow, Journal of Dairy Science 87 (2004) 2254–2267. 10.3168/jds.S0022-0302(04)70046-6.

[13] M.R. Borchers, Y.M. Chang, I.C. Tsai, B.A. Wadsworth, J.M. Bewley, A validation of technologies monitoring dairy cow feeding, ruminating, and lying behaviors, Journal of Dairy Science 99 (2016) 7458–7466. 10.3168/jds.2015-10843.

[14] P. Guarnido-Lopez, Y. Pi, J. Tao, E.D.M. Mendes, L.O. Tedeschi, Computer vision algorithms to help decision-making in cattle production, Animal Frontiers 14 (2024) 11–22. 10.1093/af/vfae028.

[15] V.M. Araújo, I. Rili, T. Gisiger, S. Gambs, E. Vasseur, M. Cellier, A.B. Diallo, AI-powered cow detection in complex farm environments, Smart Agricultural Technology 10 (2025) 100770. 10.1016/j.atech.2025.100770.

[16] Y. Zhang, P. Sun, Y. Jiang, D. Yu, F. Weng, Z. Yuan, P. Luo, W. Liu, X. Wang, ByteTrack: Multi-object Tracking by Associating Every Detection Box, in: S. Avidan, G. Brostow, M. Cissé, G.M. Farinella, T. Hassner (Eds.), Computer Vision – ECCV 2022, Springer Nature Switzerland, Cham, 2022: pp. 1–21. 10.1007/978-3-031-20047-2_1.

[17] G. Bertasius, H. Wang, L. Torresani, Is Space-Time Attention All You Need for Video Understanding?, (2021). 10.48550/arXiv.2102.05095.

[18] C. Feichtenhofer, H. Fan, J. Malik, K. He, SlowFast Networks for Video Recognition, in: 2019 IEEE/CVF International Conference on Computer Vision (ICCV), 2019: pp. 6201–6210. 10.1109/ICCV.2019.00630.

[19] S. Rao, E. Garcia, S. Neethirajan, Video-based cattle behaviour detection for digital twin development in precision dairy systems, Npj Vet. Sci. 1 (2026) 3. 10.1038/s44433-026-00004-x.

[20] T.P. Tylutki, D.G. Fox, V.M. Durbal, L.O. Tedeschi, J.B. Russell, M.E. Van Amburgh, T.R. Overton, L.E. Chase, A.N. Pell, Cornell Net Carbohydrate and Protein System: A model for precision feeding of dairy cattle, Animal Feed Science and Technology 143 (2008) 174–202. 10.1016/j.anifeedsci.2007.05.010.

[21] National Academies of Sciences, Engineering, and Medicine; Division on Earth and Life Studies; Board on Agriculture and Natural Resources; Committee on Nutrient Requirements of Dairy Cattle, Nutrient Requirements of Dairy Cattle: Eighth Revised Edition, National Academies Press (US), Washington (DC), 2021. http://www.ncbi.nlm.nih.gov/books/NBK600603/ (accessed April 26, 2026).

[22] W. Kritzinger, M. Karner, G. Traar, J. Henjes, W. Sihn, Digital Twin in manufacturing: A categorical literature review and classification, IFAC-PapersOnLine 51 (2018) 1016–1022. 10.1016/j.ifacol.2018.08.474.

[23] D. Jones, C. Snider, A. Nassehi, J. Yon, B. Hicks, Characterising the Digital Twin: A systematic literature review, CIRP Journal of Manufacturing Science and Technology 29 (2020) 36–52. 10.1016/j.cirpj.2020.02.002.

[24] A. Fuller, Z. Fan, C. Day, C. Barlow, Digital Twin: Enabling Technologies, Challenges and Open Research, IEEE Access 8 (2020) 108952–108971. 10.1109/ACCESS.2020.2998358.

[25] S. Neethirajan, Digital Twins for Cows and Chickens: From Hype Cycles to Hard Evidence in Precision Livestock Farming, Agriculture 16 (2026) 166. 10.3390/agriculture16020166.

[26] S. Neethirajan, B. Kemp, Digital Twins in Livestock Farming, Animals 11 (2021). 10.3390/ani11041008.

[27] S. Rao, S. Neethirajan, Computational Architectures for Precision Dairy Nutrition Digital Twins: A Technical Review and Implementation Framework, Sensors 25 (2025) 4899. 10.3390/s25164899.

[28] S. Rao, S. Neethirajan, Real-time identity-preserving dairy cow activity recognition for nutritional digital twin inference, [conference poster], presented at 39th IEEE Canadian Conference on Electrical and Computer Engineering (CCECE 2026), Montreal, Canada, 2026. 10.13140/RG.2.2.28915.08481.

[29] I. Halachmi, Y. Ben Meir, J. Miron, E. Maltz, Feeding behavior improves prediction of dairy cow voluntary feed intake but cannot serve as the sole indicator, Animal 10 (2016) 1501–1506. 10.1017/S1751731115001809.

[30] X.-L. Wu, K.L. Parker Gaddis, J. Burchard, H.D. Norman, E. Nicolazzi, E.E. Connor, J.B. Cole, J. Durr, An alternative interpretation of residual feed intake by phenotypic recursive relationships in dairy cattle, JDS Communications 2 (2021) 371–375. 10.3168/jdsc.2021-0080.

[31] E. Hurtado, C. Suryadevara, J. Walker, G. Tayag, H. Ryait, Digital twins as decision-support tools for automation in agriculture: A case study on robotic vaccination, Smart Agricultural Technology 13 (2026) 101675. 10.1016/j.atech.2025.101675.

[32] Y. Yamamoto, K. Akizawa, S. Aou, Y. Taniguchi, Entire-barn dairy cow tracking framework for multi-camera systems, Computers and Electronics in Agriculture 229 (2025) 109668. 10.1016/j.compag.2024.109668.

[33] H.-S. Sim, H.-C. Cho, Enhanced DeepSORT and StrongSORT for Multicattle Tracking With Optimized Detection and Re-Identification, IEEE Access 13 (2025) 19353–19364. 10.1109/ACCESS.2025.3535092.

[34] K. Abbas, Z. Afzal, A. Raza, T. Mansouri, A.W. Dowsey, C. Inchaisri, A. Alameer, Vision transformer-based multi-camera multi-object tracking framework for dairy cow monitoring, Smart Agricultural Technology 12 (2025) 101525. 10.1016/j.atech.2025.101525.

[35] M. Saar, Y. Edan, A. Godo, J. Lepar, Y. Parmet, I. Halachmi, A machine vision system to predict individual cow feed intake of different feeds in a cowshed, Animal 16 (2022) 100432. 10.1016/j.animal.2021.100432.

